# BBIBP-CorV (Sinopharm) vaccination- induced immunity is affected by age, gender and prior COVID-19 and activates responses to spike and other antigens

**DOI:** 10.1101/2022.11.30.518633

**Authors:** Zahra Hasan, Kiran Iqbal Masood, Shama Qaiser, Erum Khan, Areeba Hussain, Zara Ghous, Unab Khan, Maliha Yameen, Imran Hassan, Muhammad Imran Nasir, Muhammad Farrukh Qazi, Haris Ali Memon, Shiza Ali, Sadaf Baloch, Zulfiqar A. Bhutta, Marc Veldhoen, J. Pedro Simas, Syed Faisal Mahmood, Rabia Hussain, Kulsoom Ghias

**Author notes:** Corresponding author:* Zahra Hasan, Professor, Department of Pathology and Laboratory Medicine, The Aga Khan University (AKU), Stadium Road, P.O. Box 3500, Karachi 74800, Pakistan.

## Abstract

Long-term solutions against SARS-CoV-2 infections require understanding of immune protection induced by different vaccine COVID-19 formulations. We investigated humoral and cellular immunity induced by Sinopharm (BBIBP-CorV) in a region of high SARS-CoV-2 seroprevalence.

Levels of IgG antibodies to SARS-CoV-2 spike protein and its receptor-binding domain (RBD) were determined 24-weeks. Cellular immunity was investigated using a commercially available IFN-γ release assay to SARS-CoV-2 spike (Ag1 and 2) and extended genome antigens (Ag3).

Increasing IgG seropositivity to Spike protein and RBD was observed post-vaccination. Seropositivity was reduced in those over 50 years and raised in females and those with prior COVID-19. After 20 weeks post-vaccination, only one third of participants had positive T cell responses to SARS-CoV-2 antigens. Prior COVID-19 impacted IFNγ responses, with reactivity enhanced in those infected earlier. The frequency of IFNγ responses was highest to extended genome antigen set.

Overall, BBIBP-CorV- induced antibody responses were impacted by age, gender and prior COVID-19. Cellular immunity was present in a limited number of individuals after 20 weeks but was enhanced by prior infection. This suggests the need for booster vaccinations in older individuals. BBIBP-CorV-induced cellular activation is broader than to spike, requiring further study to understand how to monitor vaccine effectiveness.

## Introduction

COVID-19 has resulted in over 6.3 million deaths globally as a consequence of more than 643 million infections (22 November 2022) (1). Vaccinations against COVID-19 were first introduced at the start of 2021 and rolled out in stages based on both access and availability in different populations. COVID-19 vaccines included formulations based on mRNA expression of spike protein, adenovirus vector-based vaccines, and inactivated vaccine types (2). They have had a major impact on controlling both morbidity and mortality from COVID-19 (3, 4), leading to what in 2022 is thought to be the tail end of the pandemic (5).

Case fatality rates (CFR) from COVID-19 have varied greatly, ranging during the first wave in March 2020 from, 6.2% in Italy and 3.6% in Iran, to 0.79% in South Korea (6, 7). However, the CFR in Pakistan did not rise above 2% (8). In a country of over 200 million, there were 30,600 deaths reported due to COVID-19 (22 November 2022) (8), with a relatively low related morbidity (9). Pakistan has a young population, with 64% of individuals aged < 30 years. (10). Younger age has been associated with favorable outcomes in COVID-19 (11). However, the reasons for differential CFRs between countries and regions are unclear but have been related to life expectancy, population pyramids, potential exposure to other coronaviruses, and prior immunity (9).

Protection against SARS-CoV-2 infection is dependent on appropriate activation of innate and adaptive immune responses. The spike protein of SARS-CoV-2 is highly immunogenic and antibodies to spike protein are associated with protective immunity against the virus (12, 13). High seropositivity rates following natural infection have been associated with high population densities and rapid transmission resulting in high rates of infection. Studies from 2020 showed antibody seroprevalence to range between 15 to 21% in some areas of Karachi, the largest urban center in Pakistan (14). By December 2020, we observed IgG antibodies to spike protein to be present in greater than 50% of unvaccinated healthy blood donors (15). By April 2021, we observed IgG antibody seroprevalence to be up to 83% in an uninfected health care cohort in Karachi (16).

Early activation of T cell producing IFN-γ and IL-2 indicates robust T cell effector response in patients with COVID-19 are associated with decreased disease severity (17). The induction of effective SARS-CoV-2-specific CD4+ T cells is also linked with effective activation of antibody responses. T cell reactivity associated with viral clearance is linked with recognition of spike and nucleocapsid proteins (18, 19).

About 12 billion doses of COVID-19 vaccinations have been administered worldwide, with 66% of the global population having received one dose (20). Most of the studies on vaccine efficacy and immunological responses are based on the analysis of populations vaccinated by mRNA type (Pfizer-BNT162b2-, Moderna mRNA-1273) and vector-based vaccines (ChAdOx nCoV-19). The vaccine, Sinopharm (inactivated) prepared by Beijing Bio-Institute of Biological Products Co-Ltd. (BBIBP) also called BBIBP-CorV (Vero cells) is an aluminum-hydroxide-adjuvanted, inactivated whole-virus vaccine. It is administered in two doses, given four weeks apart (21). 1.65 billion doses of Sinopharm have been delivered to 79 countries worldwide (22 November 2022) (22). There is limited data on inactivated vaccines such as those administered in many low-middle income countries (23, 24).

In Pakistan, 139 million individuals (70%) of the population have received their first dose of vaccination, with 132 million (66%) individuals fully vaccinated (22 November 2022) (8). Vaccinations were rolled out in February 2021 as emergency use authorization (EUA) was obtained by BBIBP-CorV as one of the primary vaccines administered (25). Vaccinations were given in a stratified manner that is first, to health care workers and then to older age groups in the country.

A recent study from Pakistan has shown that full vaccination with BBIBP-CorV was effective in reducing the risk of symptomatic COVID-19 infection (94.3%), hospitalisations (60.5%) and mortality (98.6%), respectively (26). Studies on immune response induced by BBIBP-CorV have shown differing results. In Sri Lanka, more than 90% seropositivity was observed 6 weeks post-vaccination (24, 27). A study from the United Arab Emirates showed seropositivity of 78% in the general population after full vaccination (28), with a similar result in a report from Pakistan (29).

Humoral responses against SARS-CoV-2 are driven both by natural infection and COVID-19 vaccinations (12, 13, 30, 31). It is particularly important to have data from high infectious disease burden regions with limited COVID-19 vaccine access where primarily inactivated vaccines were administered. In this study, we investigated the dynamics of BBIBP-CorV-induced IgG responses to spike and receptor binding domain (RBD) proteins for up to 25 weeks post-vaccination in a healthcare associated cohort in Karachi, Pakistan (32). We investigated humoral and cellular immunity to SARS-CoV-2 antigens to determine how these are impacted by age, sex, and prior history of COVID-19 infection.

## Methods

This study was approved by the Ethical Review Committee of Aga Khan University (AKU) (project #2020-5152-11688).

### Study subjects

Vaccinations were administered from February 2021 at AKU Hospital, in a designated COVID-19 Vaccination Center by the Department of Health, Government of Sindh. BBIBP-CorV vaccination was administered as per guidelines of the National Covid Operation and Command (NCOC), Government of Pakistan (8). The vaccine route was an intramuscular injection in the deltoid area. The time interval between the first and second doses of Sinopharm/BBIBP-CorV was four weeks as per manufacturer’s recommendations (21).

We recruited adult study subjects aged over 18 years with written informed consent after they had received their first dose of BBIPP-CorV vaccination. Subjects were recruited by a consecutive convenience sampling method. They included healthcare workers, Aga Khan University (AKU) employees, and their family members who volunteered to participate in the study. A verbal history of prior COVID-19 infection was taken at the time of recruitment. A SARS-CoV-2 PCR test was used to identify laboratory confirmed COVID-19 in all cases. Positive antibody results if available were also documented. There was no bias in selection of cases based on any prior COVID-19 history. During the study period, active surveillance of infection was not performed. However, subjects were encouraged to inform the study team if they had PCR-confirmed COVID-19 infection during the post-vaccination study period.

AKUH provided SARS-CoV-2 PCR testing to all employees through the pandemic. Therefore, individuals with symptoms associated with COVID-19 and those who had exposure to laboratory confirmed COVID-19 cases were able to get themselves tested free of cost.

In total, we enrolled 312 participants to the study. All of them were requested to give blood samples for up to four time points during the study period, 24 weeks after their first dose of BBIBP-CorV (Figure 1). The number of samples given by participants varied as some refused subsequent blood draws. In all, 312 participants submitted ≥ 1 test; 248 underwent ≥ 2 tests, 151 gave ≥ 3 samples and 41 gave 4 samples (Figure 1).

**Figure 1.**
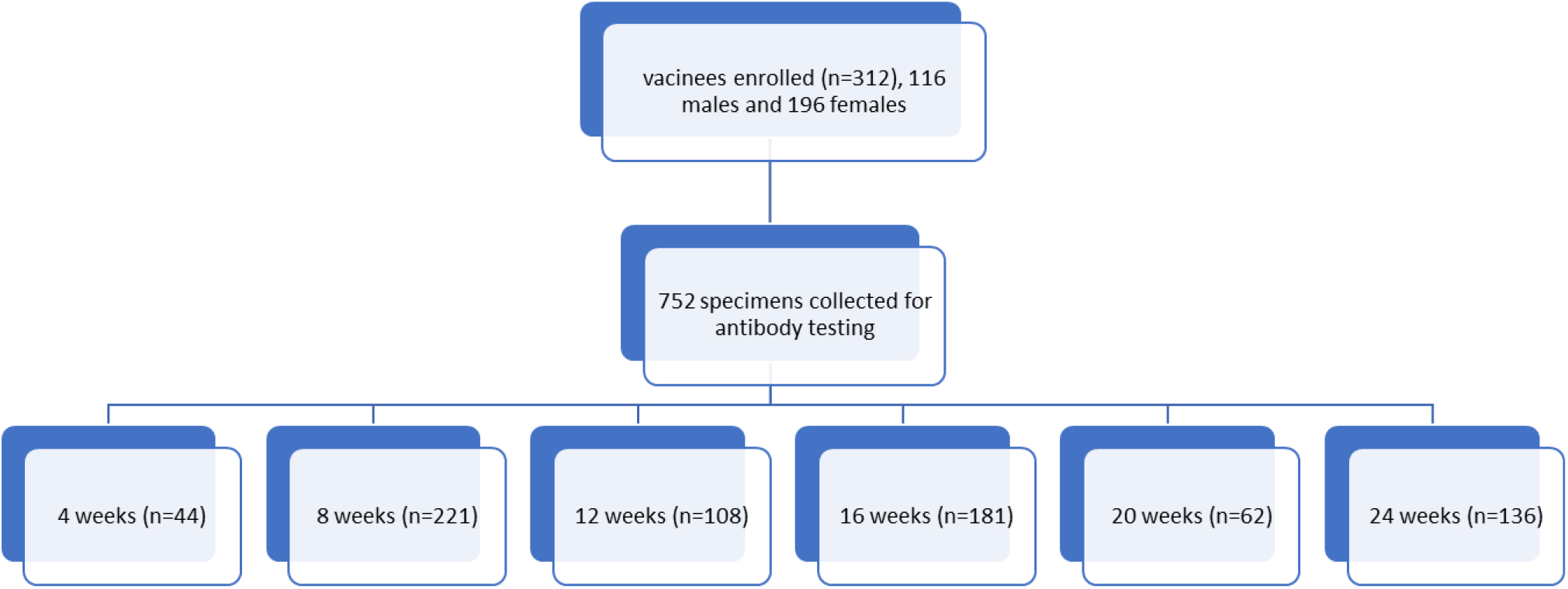
Overview of BBIBP-CorV vaccination study plan

### Sample collection

Three milliliters of whole blood was collected in a serum collection tube. Serum was separated from blood and stored at −80°C. All sera were inactivated at 56°C for 30 min before testing.

### Recombinant Proteins

Recombinant Spike and RBD proteins were produced by the laboratory of Prof. Paul Alves, iBET, NOVA ITQB University, Portugal. The proteins were extensively characterized and found to be both stable and consistent for use in serological assays (33, 34).

### ELISA for IgG to Spike and RBD

The assay used was based on the protocol developed by the laboratory of Prof. Florian Krammer (35, 36) and received FDA authorization (37). All serum samples were diluted at 1:100 and tested in duplicate using an in-house enzyme-linked absorbent assay (ELISA) (36) and as per the protocol described by Figueiredo-Campos *et al*. (13).

This assay has been validated in our laboratory using sera from healthy individuals from the pre-pandemic period as negative controls (38).The cut-off was established by calculating the Mean +3SD (OD 0.5 at 450nm) for IgG measurements of Spike and RBD in the pre-pandemic controls.

Duplicate samples of pooled positive IgG sera at each dilution were assessed to check for intra- and inter-plate variability (Supplementary Figs. 1-2). The sensitivity and specificity of the ELISA assays for IgG to spike protein was found to be 100% (92.1-100, 95% CI) with a specificity of 100% (93.5-100, 95% CI). The ELISA for IgG to RBD had a sensitivity of 91.1% (78.8-97.5, 95% CI) with a specificity of 94.6% (82.4-98, 95% CI) (15).

Briefly, SARS-CoV-2 Spike and/or RBD protein were used to coat plates with 50μl of Spike or RBD protein at a concentration of 2 μg/ml in PBS. Wells were blocked and then incubated with 100 μl serum samples for 2 hours. Wells were washed and stained with secondary antibody conjugated with Horse Radish Peroxidase (HRP). Plates were read for color intensity as optical density units (OD) at 450nm using an Elisa reader.

### Statistical analysis

The Statistical Package for the Social Sciences version 24.0 (SPSS Inc. Chicago, IL, USA, 2013) was used to carry out descriptive statistics of participants for demographic variables (gender, age). In addition, a history of previous COVID-19 infection was analysed. The normality of data was checked through Shapiro-Wilk test; mean ±SD was used to describe normally distributed and median (IQR) for skewed continuous data. Data are presented in both frequencies and percentages. Chi-square test was used to compare the frequencies of IgG antibodies with respect to age groups (either less than 50 years or those 50 years and above), gender and COVID-19 infection as variables. The threshold of significance was a p-value <0.05.

A multivariable analysis was run to have age, gender and prior history of COVID-19 adjusted to the frequency of positive IgG responses determined in each condition. An odds ratio was used to determine significance together with a p value <0.05.

Correlation between the RBD titers and neutralizing potential was determined using the Spearman’s rank correlation test using the GraphPad prism. A p value <0.05 was considered as significant.

### QuantiFERON SARS-CoV-2

Five ml of whole blood was collected in a lithium heparin collection tube. QuantiFERON ELISA (Cat. No. 626410) and QuantiFERON SARS-CoV2 RUO Starter + Extended Pack (Cat. No. 626915); QIAGEN, Hilden, Germany was used as the interferon gamma release assay (IGRA). This assay consists of three Antigen tubes, SARS-CoV-2 Ag1, Ag2 and Ag3, that use a combination of proprietary antigen peptides specific to SARS-CoV-2 to stimulate lymphocytes involved in cell-mediated immunity in heparinized whole blood. The QFN SARS-CoV-2 Ag1 tube contains CD4 + epitopes derived from the S1 subunit (Receptor Binding Domain) of the Spike protein, the Ag2 tube contains CD4 +CD8 epitopes from the S1 and S2 subunits of the Spike protein and the Ag3 tube consists of CD4 + CD8 + epitopes from S1 and S2, and epitopes from M and rest of the genome.

Plasma from stimulated samples were used for detection of IFN-γ using an enzyme-linked immunosorbent assay (ELISA)-based platform. Following ELISA, quantitative results (IFN-γ concentration in IU/ml) were recorded and used for analysis. Elevated response was defined as a value at least 0.15 IU/mL greater than the background IU/mL value from the QFN-SARS-CoV-2 Nil tube.

## Results

### Description of study

Our study participants received their first dose of BBIBP-CorV vaccine between February and June 2021. Blood samples were collected at 4, 8, 12, 16, 20 and 24 weeks post-vaccination (Figure 1). The age range of study subjects was 20 to 101 years with a mean age of 40.7± 16.5 years (Table 1). Seventy four percent of subjects (n=231) were aged 50 years or lower, 26% (n=81) greater than 50 years; while 63% were females.

**Table 1.** Characteristics of study subjects.

| Group | n (%) | Females | Males | p value |
| --- | --- | --- | --- | --- |
| Total | 312 (100) | 196 (62.8%) | 116 (37.2%) | ns |
| ≤ 50 years (n, %) | 231 (74) | 154 (66.7) | 77 (33.3) | ns |
| >50 years (n, %) | 81 (26) | 42 (52) | 39 (48) | ns |
| History of COVID-19 |  |  |  |  |
| Yes (n, %) | 89 (28.8%) | 60 (67.4) | 29 (32.6) | ns |
| **Age** | **(years Mean, SD)** |  |  |  |
| **Total** | **44 (14)** |  |  |  |
| COVID-19 pre vaccination | 34.4 (10.9) |  |  | 0.0039 |
| COVID-19 post vaccination | 44.2 (13.8) |  |  |  |
| **Total with prior COVID-19 (n, %)** | **68 (21.8%)** | **47 (69.1)** | **21 (30.9)** | **ns** |
| Age-wise |  |  |  | ns |
| ≤ 50 years |  | 40 (85.1) | 19 (90.5) | ns |
| >50 years |  | 7 (14.9) | 2 (9.5) | ns |
| All post-vaccination COVID-19 | 21 (7.1%) | 13 (62) | 8 (38) | ns |
| Age-wise |  |  |  | ns |
| ≤ 50 years |  | 4 | 9 | ns |
| >50 years |  | 4 (30.8) | 4 (50) | ns |

Eighty-nine individuals (29%) had COVID-19. Of these, 68 (22%) had COVID-19 prior to enrollment in the study; median period 23 weeks prior to enrollment (range 3 – 56 weeks). 21 individuals (7%) reported COVID-19 for the first time after full vaccination.

Of note, the age of individuals who had COVID-19 before the study was significantly lower than those who got COVID-19 post-vaccination (p=0.0039). We did not find any difference between sex and the age groups of individuals between those who had a prior history of COVID-19 as compared with those who were infected after enrolment in the study.

#### Dynamics of IgG Antibody responses to Spike protein after BBIBP-CorV vaccination

IgG antibody levels to Spike protein were measured in individuals tested between 4 and 24 weeks after their first dose of BBIBP-CorV vaccination. IgG levels increased post-vaccination, p<0.0001, Kruskal-Wallis test (Fig. 2A). However, it was observed that the magnitude of IgG antibodies was lowered at 16 as compared with 8 weeks post-vaccination (p<0.0001) (Fig. 2A). No difference in IgG levels was observed between 20- and 24-weeks post-vaccination. By 4 weeks after vaccination, 57% of individuals were seropositive, increasing to 87% at 8 weeks. Seropositivity was 82% at 20 weeks and 90% by 24 weeks (Fig. 2B).

**Figure 2.**
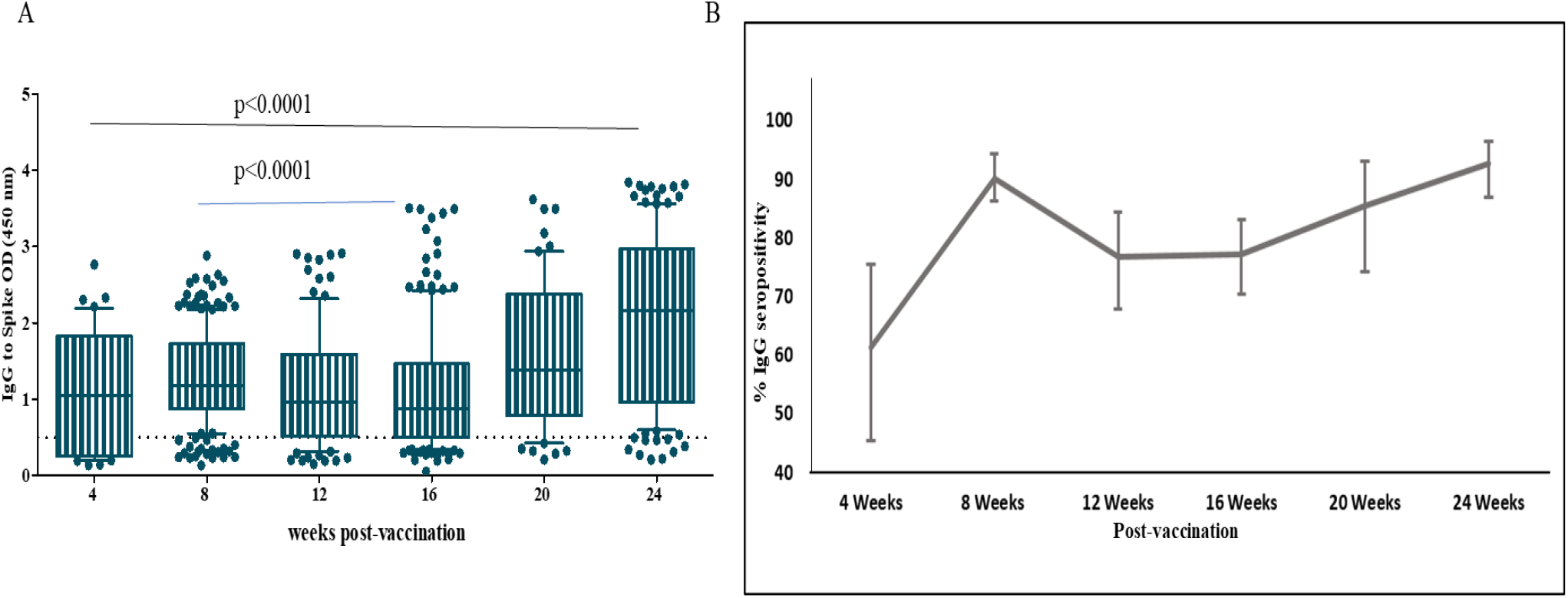
Dynamics of IgG antibody levels to Spike protein after BBIBP-CorV vaccination. IgG antibodies were determined in sera of individuals after 4, 8, 12, 16, 20 and 24 weeks, postvaccination. Graphs show (A) IgG levels, cut-off for positive responses at 0.5 OD 450 nm is indicated by a dotted horizontal line. Data is depicted as a box and whiskers plot with IQR (10-90th) percentile indicated by error bars. A horizontal line represents median values. (B) the frequency of IgG seropositive individuals at different time intervals is shown.

#### Dynamics of IgG antibodies to RBD after BBIBP-CorV vaccination

Although RBD is an integral part of Spike protein, we also determined IgG antibody levels to RBD separately, as IgG to this peptide is associated with neutralizing antibody to SARS-CoV-2 and hence considered protective. Study participants were tested between 4 and 24 weeks after their first dose of vaccination. IgG levels were found to increase significantly across the study period, p<0.0001 (Fig. 3A). IgG antibody levels were higher at −16 (p=0.036), −20 (p<0.0002), and - 24 weeks (p<0.0001) as compared with those at 8 weeks post-vaccination. No difference was observed between IgG titers to RBD between 20 and 24 weeks.

**Fig. 3.**
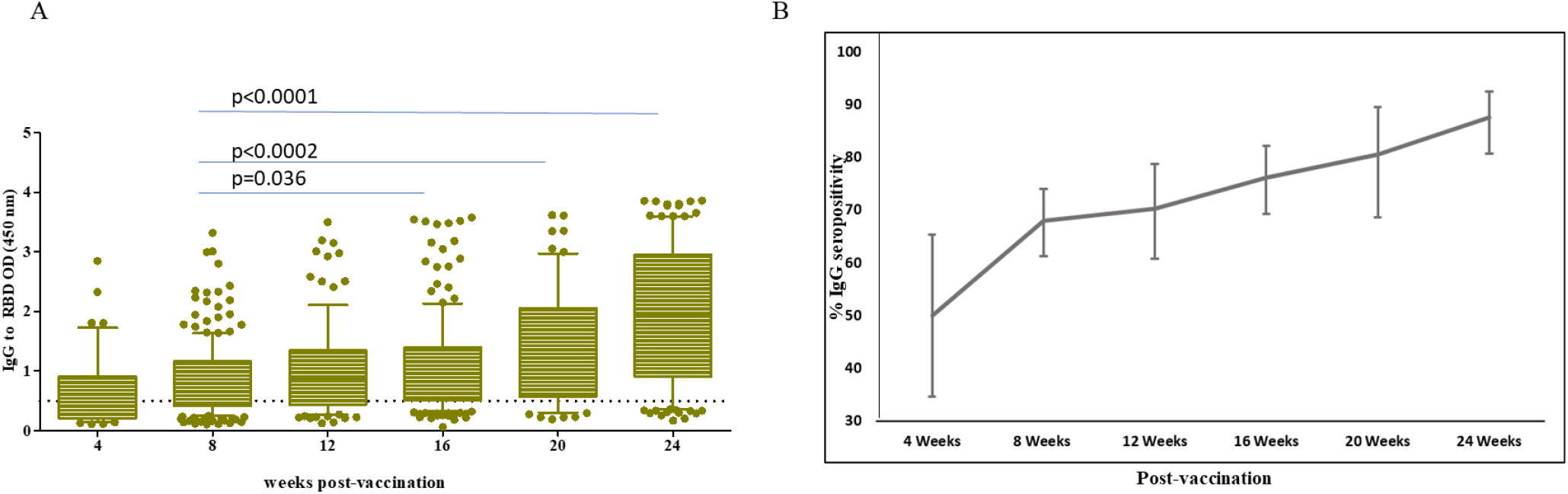
Dynamics of IgG antibody levels to RBD protein and after BBIBP-CorV vaccination. IgG antibodies were determined in sera of individuals after 4, 8, 12, 16, 20 and 24 weeks, post-vaccination. Graphs show (A) IgG levels, cut-off for positive responses at 0.5 OD 450 nm is indicated by a dotted horizontal line. Data is depicted as a box and whiskers plot with IQR (10-90^th^) percentile indicated by error bars. A horizontal line represents median values.(B)the frequency of IgG seropositive individuals at different time intervals is shown.

After 4 weeks of receiving their first dose of BBIBP-CorV, 48% of individuals showed seropositivity to RBD. This increased further between 8 and 24 weeks; 61%, 62 %, 68%, 73% and 85 % at −8, −12, −16, - 20 and 24 weeks respectively (Fig. 3B).

#### Relationship between IgG antibody levels to Spike protein and RBD

We further compared BBIBP-CorV vaccination-induced IgG responses to Spike and RBD by comparing IgG levels. The magnitude of antibodies to Spike was greater compared to RBD at 4 and 8 weeks post-vaccination (MWU; p<0.024, p<0.001, respectively, Supplementary figure 3A-B). At subsequent time intervals, there was no difference found between antibody levels to Spike and RBD (Supplementary figure 3C-F).

We further analyzed concordance of IgG antibodies to Spike and RBD. Spearman’s rank correlation test was applied to assess the significance of concordance between IgG levels to Spike and RBD at each time intervals (−4, −8, −12, −16, −20 and −24 weeks post vaccination; Supplementary figure 4).

#### Higher IgG seropositivity in younger individuals and in females

We used a multi-variate model to investigate the impact of age and sex on IgG antibody responses after BBIBP-CorV vaccination. The frequency of seropositive individuals present at each time interval was stratified by age (<50 >50 years), gender and a prior history of COVID-19 to adjust for these variables in analyzing the results of the IgG antibody responses to Spike protein and RBD.

We observed that IgG seropositivity to Spike was greater in those aged <50 years compared with other individuals >50 years, at −8 (p<0.001), −12 (p<0.001), and −16 (p=0.015) weeks post vaccination (Fig. 4A). In the case of IgG anti RBD antibodies, the frequency of seropositive individuals was greater in those < 50 years at 8 weeks post vaccination, p=0.003 (Fig. 4B).

**Figure 4.**
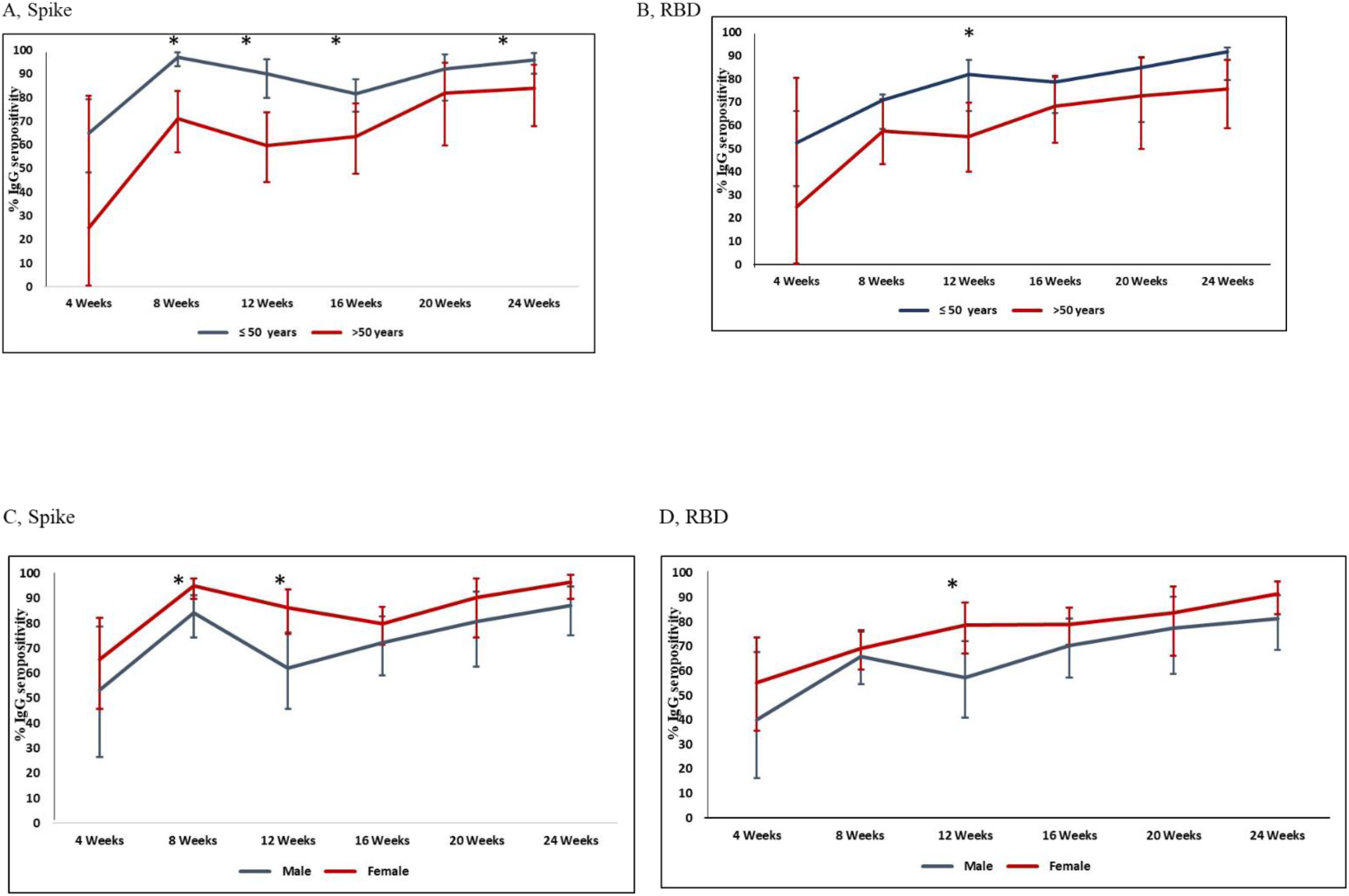
Effect of age and gender on IgG responses to Spike and RBD after BBIBP-CorV vaccination. Graphs depict the frequency of IgG seropositivity to individuals tested (x axis) −4, −8, 12, −16, 20 and 24 weeks, post-vaccination. Graphs compare seropositivity between individuals aged 50 years and below to those aged above 50 years in response A) Spike and B) RBD. Seropositivity of IgG responses compared based on gender is shown to C)Spike and D) RBD. ‘*’ indicate p values ≤ 0.05 at each particular time point between groups. The data is indicated as 95% CIs with error bars at +-2SD.

We also observed an effect of gender and that IgG seropositivity was higher amongst females particularly in the context of IgG antibodies to Spike at −8 (p=0.01) and −12 (p=0.005) weeks post vaccination (Fig. 4C). IgG seropositivity to RBD was also higher in females than in males at 12 weeks (p=0.018) post-vaccination (Fig. 4D).

#### Prior COVID-19 enhances BBIBP-CorV-induced IgG responses

We also investigated the impact of prior COVID-19, on IgG antibody responses to Spike and RBD post-vaccination. Of our participants, 89 (29%) had confirmed COVID-19, either prior to (n=68) or after vaccination (n=21). We first focused on those who had COVID-19 prior to enrollment; 63 (92.6%) individuals were < 50 years and 5 (7.4%) were > 50 years. All participants with prior COVID-19 had a positive IgG response to spike and RBD at the time they were enrolled in this study. None of them developed COVID-19 a second time after vaccination and during the follow-up period of the study. IgG seropositivity to Spike was greater after 16 weeks (p=0.013) as compared with those without prior infection (Fig. 5A). IgG seropositivity to RBD was greater after 8 weeks (p=0.012) in those who suffered COVID-19 (Fig. 5B), Supplementary Table 4.

**Fig. 5.**
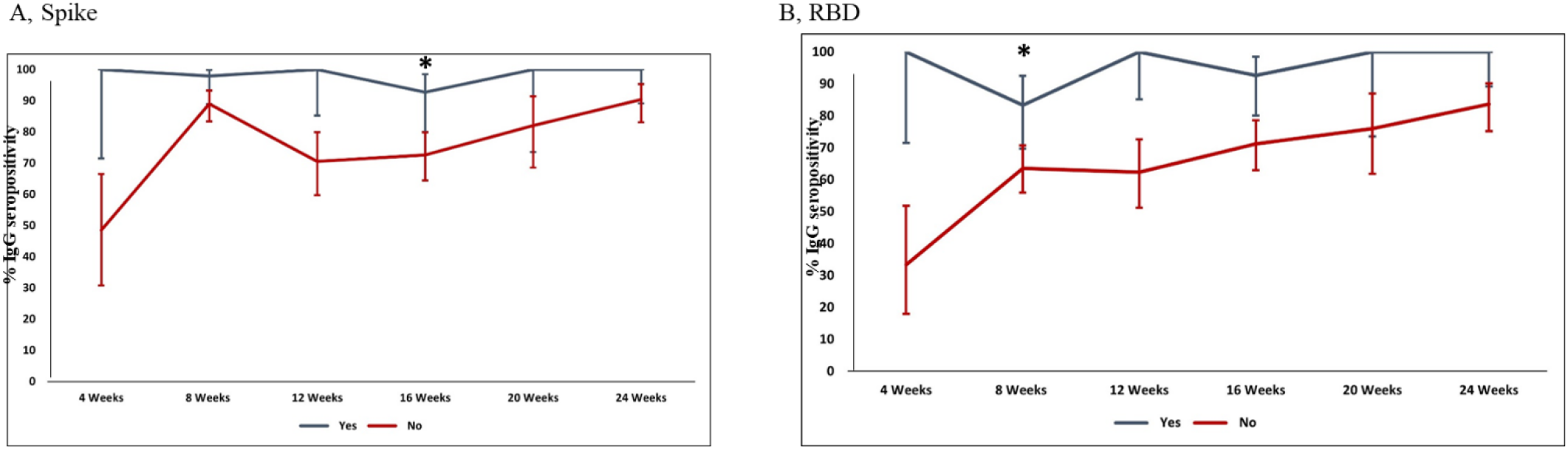
Effect of prior COVID on IgG responses to Spike and RBD after BBIBP-CorV vaccination. Graphs depict the frequency of IgG seropositivity in individuals tested (x axis) −4, −8, 12, −16, 20 and 24 weeks after vaccination. Graphs compare seropositivity between individuals who had COVID-19 prior to vaccination as compared with those who did not have COVID-19. IgG responses are shown to A) Spike and B) RBD. ‘*’ indicate p values ≤ 0.05 between groups. The data is indicated as 95% CIs with error bars at +-2SD.

In the 21 participants who developed COVID-19 after vaccination, the average time interval of acquiring infection was 16 weeks after the first dose of BBIBP-CorV vaccine. Thirteen (62%) participants were < 50 years, with 8 (18%) > 50 years. of those who got COVID-19 after enrollment, 52% were initially negative for IgG to RBD; measured at 4 weeks (n=2) and 8 weeks (n= 9). All of these developed IgG antibodies to RBD when tested after their COVID-19 diagnosis.

### SARS-CoV-2 antigen induced cellular reactivity

We next analyzed cellular immunity after BBIBP-CorV vaccination, using a whole blood assay on samples drawn between 20 and 24 weeks post-vaccination. This period was coincident with a maximal IgG antibody response to both Spike and RBD and importantly at this time we had not observed any impact of age, gender or prior COVID-19 on seropositivity to Spike or RBD (Figs 4-5). IFNγresponses in whole blood samples stimulated with Ag1 tube contains CD4 + epitopes derived from the S1 subunit (RBD), the Ag2 tube contains CD4 +CD8 epitopes from the S1 and S2 subunits of the Spike protein and the Ag3 tube consists of CD4 + CD8 + epitopes from S1 and S2, and also epitopes from M and rest of the genome.

Participants had a mean of 45 years SD 16 years (range 19 – 82 years). Sixty-eight (68.7%) percent were aged below or equal to 50 years whilst 31 (31.3%) were aged 50 years and above. There were 41 (41.4%) males and 58 (58.6%) females. 15 had a prior history of COVID-19 disease (67 % females and 33% males), Table 2. When T cell responses were stratified according to different antigens; IFNγ stimulation was observed in response to each, Ag1 (p=0.003), Ag2 (p<0.001) and Ag3 (p<0.001), (Supplementary figure 5A, Supplementary Table 2). Overall, 37.4% of donors had IFN-γ producing T cells to one or more antigens (Table 2). According to different antigens, positive IFNγ responses were seen specifically in; 16% to Ag1, 24% to Ag2 and 33% to Ag3, Supplementary figure 5B.

**Table 2.** Individuals with prior COVID-19 show increased IFN-g responses to SARS-CoV-2 antigens.

| **Parameters** | *Total (n)* | *COVID-IGRA positive (n)* | *COVID-IGRA negative (n)* | *% IGRA positive* |
| --- | --- | --- | --- | --- |
|  | 99 | 37 | 62 | 37.4 |
| **Age (years)** |  |  |  |  |
| ≤ 50 | 68 | 30 | 38 | 44.1 |
| >50 | 31 | 7 | 24 | 22.5 |
| **Sex** |  |  |  |  |
| Male | 41 | 16 | 25 | 39 |
| Female | 58 | 21 | 37 | 36.2 |
| **History** |  |  |  |  |
| Prior COVID-19 | 15 | 10 | 5 | 66.6 |
| No COVID-19 | 84 | 27 | 57 | 32.1 |

Whole blood samples were tested for reactivity to antigens in the COVID-IGRA assay. Results are shown as per a positive cut of 0.15 IU/ml IFN-g as per manufacturer's instructions

We investigated IFN-γ activation whether there was an impact of age and gender of the donors (Table 2). For age-wise analysis, IFNγ responsiveness to each antigen was compared between those < 50 years, with those > 50 years. Reactivity to SARS-CoV-2 antigens was 44% in the younger age group as compared with 22.5% in the older age group. No difference was seen between cellular reactivity to Ag1, Ag2 or Ag3 between males and females.

Next, we evaluated whether having COVID-19 prior to vaccination impacted on T cell IFN-γ responses. We compared those who had COVID-19 prior to vaccination (n=15) with those who had been vaccinated but did not have a history of disease (N=84). Overall, when reactivity to either one or more antigens (Ag1-3) were considered, 66% of those who had prior COVID-19 had a positive T cell response compared to 32.5% in those uninfected. Further, we compared the effect of prior COVID-19 on cellular reactivity to different specific antigens. No difference was observed between antigen-stimulated IFN-γ levels between the two groups. However, the proportion of individuals with reactive T cells was greater in those with COVID-19 history in the case of response to Ag2 (47%in COVID, 20% in no-COVID) and Ag3 (60% in COVID, 29% in no-COVID) between groups (Fig. 6A-B).

**Figure 6.**
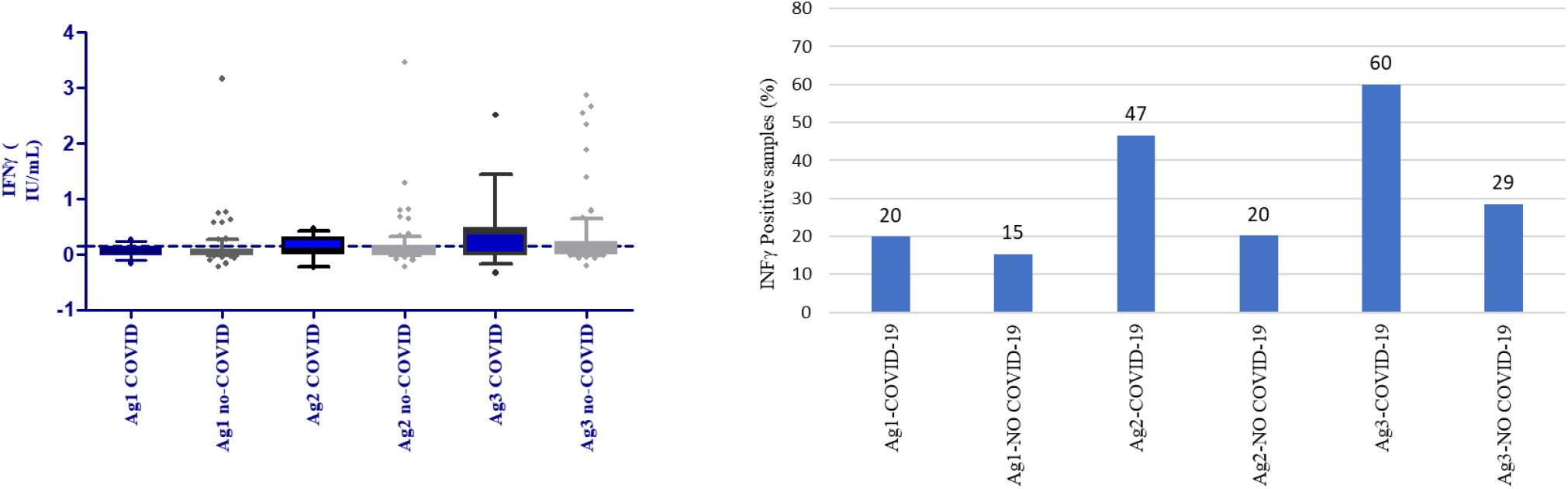
Effect of prior COVID-19 on the cellular response after BBIBP-CorV vaccination. IFN-g was measured in stimulated whole blood supernatants of donors 20-24 weeks post-vaccination using the QFN SARS CoV-2 SARS-CoV-2 assay, Ag1 tube contains CD4 + epitopes derived from the S1 subunit, Ag2 tube contains CD4 +CD8 epitopes from the S1 and S2 subunits and Ag3 tube consists of CD4 + CD8 + epitopes from S1 and S2, and also epitopes from M and rest of the genome. A, Graphs show responses to with prior-COVID (n=15) or COVID-naïve (no-COVID, n=84) individuals. IFN-g secretion is shown in response to antigen stimulation for either group. B, percentage of individuals with reactive T cells to SARS-CoV-2 antigens in each group.

### Correlation between IgG antibody and IFNγ responsiveness to SARS-CoV-2 antigens

We investigated whether there was a correlation between levels of IgG antibodies to Spike and RBD proteins, and the IFNγ secretion observed in the individuals in response to stimulation by the different antigen sets using the Quantiferon whole blood assay. Results of individuals were grouped into those who had COVID-19 prior to vaccination and those who did not have COVID-19. In the COVID-19 group, a correlation between IgG antibodies to Spike and IFNγ secretion to Ag2 (S1 and S2 epitopes) were observed, Table 2. This was absent in the case of Ag1 and Ag2. Further, no correlation was seen between IgG to RBD and IFNγ secretion of Ag1,2 or 3 tubes.

However, in individuals without prior history of COVID-19 there was a correlation observed between IgG antibodies to Spike and IFNγ responses to SARS-CoV-2 antigens for all three tube sets. The same was observed in the case of IgG antibodies to RBD and IFNγ responses to Ag1, Ag2 and Ag3 tubes.

## Discussion

Our current study provides insights into the dynamics of humoral and cellular responses post BIBB CorV vaccines. Such studies are critical to evaluating vaccine efficacy, identifying high risk groups and necessity for spacing and number of boosters required for optimal impact. Humoral responses focused on IgG antibody responses to spike protein and RBD while T cell stimulated IFN γ responses were analyzed to CD4 and CD8 epitopes from Spike and other proteins. We show that cellular responses to SARS-CoV-2 antigens were observed in only one third of individuals without a history of COVID-19 infection. Notably, antigen-stimulated IFN-γ secretion was evident to - other than spike protein, demonstrating the effect of inactivated vaccines.

BBIBP-CorV vaccinations were rolled out in February 2021 (25) first in healthcare workers and then elderly in the Pakistani population. Our study subjects were healthy adults without any significant co-morbid conditions. We observed a significant trend towards increased IgG antibodies to Spike and RBD upto 24 weeks post-vaccination with BBIBP-CorV. Notably, the seropositivity levels were slightly higher to Spike protein compared to RBD. IgG to spike and RBD showed a significant positive correlation at each of the time intervals. We found that the relative levels of IgG antibodies were higher to spike as compared to RBD at both 4- and 8-weeks post-vaccination. Differences in antibody responses to Spike and RBD are likely reflective of the increased recognition of the number of epitopes present in Spike protein as compared with its RBD sub-unit, leading to differential recognition of epitopes. The spike protein of SARS-CoV-2 shares sequence homology with other coronaviruses as well as some of the proteins in commensal gut microbiota of humans and prime immune responses (39). Antibodies to spike protein are also likely due to cross-reactive responses to other pathogens such as, human coronaviruses (40). We have previously observed higher titers of IgG Spike as compared with RBD in healthy uninfected individuals (38). Crossreactivity to Spike has been demonstrated in the context of antibody responses to full-length spike proteins using a peptide microarray system (41, 42).

Full vaccination period is considered as 2 weeks or more after the second dose of a two-dose vaccine regimen. We found seropositivity to be 87.3% to Spike and 61.5% to RBD at 8 weeks post-vaccination. Our data differ from those reported in a study from Sri Lanka by Jeewandara *et al*. of BBIBP-CorV vaccinated group at 0, 4 and 6 weeks, which reported 95% seroconversion at 6 weeks and also found these associated with neutralizing activity to SARS-CoV-2 (43). However, it correlates with the lower (43) antibody responses to BBIBP-CorV measured in dialysis patients in the UAE measured at 6 weeks of vaccination (28). Also, with data from Pakistan that showed 78% of antibody responses to RBD in individuals vaccinated with BBIBP-CorV (29).

BBIBP-CorV vaccination-induced IgG seropositivity to spike protein followed the same trend in those aged below 50 years as compared with those aged 50 years and above but seropositivity were significantly higher in the younger age group. Similarly, younger individuals had higher seropositivity to RBD than those who were older. These data correlate with those reported by Ferenci *et al*. from Hungary who showed that RBD-specific antibody responses after 2 doses of BBIBP-CorV were present in 90% of cases below 50 years but were reduced in those who were older (44). BBIBP-CorV vaccination data from Sri Lanka also shows reduced immune responses in individuals aged 60 years and above (43). Such finding warrant a relook at boosters among different age group.

IgG to RBD is associated with neutralizing activity (45) (46). We found IgG to RBD measured in both COVID-19 convalescent and healthy individuals to be associated with neutralizing activity to SARS-CoV-2 (47). Aijaz et al. reported from Pakistan that antibodies to RBD after BBIBP-CorV vaccination were associated with neutralizing antibodies (29). BNT162b2 mRNA and ChAdOx vaccinations have been shown to effectively induce IgG antibodies to spike and neutralizing antibodies, within 14 days of the second dose, or full vaccination (45, 48). The slower rise in BBIBP-CorV-induced IgG to RBD we observed correlates with earlier reports (24).

A possible explanation for the difference observed between early post-vaccination responses between younger and older age groups may be related to T independent B cell expansion (TI). T cell activation may occur earlier in the younger age group (<50 years). T follicular helper cell independent expansion has been shown to occur in response to SARS-CoV-2 infection in mice, resulting in high affinity antibodies (49). Therefore, resident T cells may drive antigen specific B cell expansion. In the older age group, T cell activation is compromised (50) and therefore, the antibody response dramatically drops with removal of antigen antibody complexes. However, the T independent responses continue to produce IgG antibodies as likely indicated by slow rise of IgG antibodies to RBD (46).

We found that seropositivity of IgG to spike and RBD in females was greater than males after vaccination. A comparison of antibody responses to ChAdOx1 has been shown to induce higher levels in females than males (48). Sex-specific differences related to COVID-19 have been observed between males and females, with increased COVID-19 morbidity in males. Inflammatory cytokine levels shown to be raised in females than males (51-53).

Those with a history of COVID-19 displayed significantly greater higher antibody seropositivity than vaccinees without infection. This correlates with reports by Aijaz et al who observed that seropositivity rates from natural infection were greater than those induced by Sinopharm vaccination (29). Antibody positivity protects against SARS-CoV-2 reinfection for at least 7 months (54). The IgG response to natural infection by SARS-CoV-2 is thought to last up to 10 months or more (13, 55). Difference in dynamics of IgG responses with and without prior COVID-19 may be due to the re-activation of T cell responses present due to natural infection (56, 57), with cellular immunity activating B cell class switching leading to increased IgG production.

Another consideration that may impact the quality of neutralizing antibodies generated by BBIBP-CorV could be the effect of inactivation on the spike protein. Cai et al. have shown that the antigenic epitopes presented by the glycoprotein are dependent on its structural integrity (58). Therefore, virus inactivated vaccines may contain proteins that do not have the RBD domain in the open conformation required to serve as effective antigenic epitopes.

Vaccine efficacy studies of inactivated types such as, BBIBP-CorV and CoronaVac conducted in China, India, and Chile (23, 59, 60), shown that these are effective against preventing hospitalisations and severe COVID-19. In Pakistan, BBIBP-CorV vaccination was shown to reduce hospitalization, mortality and symptomatic COVID-19 in the older age group however, this was based on testing after 14 days of the second dose, or 6 weeks post-vaccination (26).

We did not observe any waning of antibodies to RBD. This differs from reports of waning humoral immune responses to BNT162b2 over 6 months in Israel demonstrated by decreased IgG levels and neutralizing antibodies over 3 months, accompanied by breakthrough infections (61). A case control study of vaccine efficacy against SARS-CoV-2 variants using BNT162b2 in Qatar also showed protective antibody levels to decrease after vaccination (62). A study in Sri Lanka showed a waning of BBIBP-CorV induced antibodies by 12 weeks after vaccination (43).

The absence of antibody waning may be a consequence of continued SARS-CoV-2 exposure in the community, boosting cross-reactive responses. The period studied was alpha and then delta variants were predominant leading to a peak in morbidity and mortality by June 2021 (63). Whilst BBIBP-CorV vaccination been shown to be effective in reducing hospitalization and symptomatic COVID-19, it is also shown that induced levels of IgG to RBD are lower than other vaccinations and therefore booster vaccinations are recommended (64, 65).

T cell immunity is associated with long term memory responses and is the hallmark of protection induced by natural infections and vaccines. The T cell responses show durable and polyfunctional virus specific memory CD4 and CD8+ T cells in infected patients up to 8 months after infection, with specificity was observed to a range of SARS-CoV-2 antigen (66).

We used a whole blood based assay to measure cellular immunity. Previous reports have shown that for measuring cellular immunity to SARS-COV-2, it is effective to detect IFN-γ to different Nucleocapsid, Membrane, Spike proteins but the spike antigen was the most efficient at detecting T cell specific response in COVID19 cases (67). The commercial assay we used has been shown to effectively determine T cell responses to BNT2162b2 mRNA vaccination responses (68). Preliminary evaluation of QuantiFERON SARS-CoV-2 (69) after mRNA vaccination showed that there was a strong cellular response is seen after the first dose, then does wane after 2 months. Enhanced cellular immunity in those with COVID infection history and vaccination has previously been shown to be the case for mRNA vaccines (70, 71). The IFN-γ release assay can be used to measures memory responses in COVID-19 convalescent individuals (72).

We show here that only one third of individuals had T cell reactivity to SARS-CoV-2 antigens 24 weeks after vaccination with BBIBP-CorV. This was doubled in individuals with a prior history of COVID-19. Earlier reports from Sri Lanka that Sinopharm vaccination elicited a strong cellular response after 4 weeks of the first dose of vaccine (27). However, there is currently no data on subsequent cellular immunity after a 6 month period.

Importantly, in our participants, we observed the strongest cellular reactivity to antigens of the extended set which included Membrane and Envelope proteins in addition to Spike antigens. This trend was evident in individuals with and without prior COVID-19. This highlights the broad nature of T cell responses to inactivated vaccines such as BBIBP-CorV.

Our study has some limitations. The vaccination rollout was rapid and on an emergency basis in our study population, healthcare worker and older age group populations. Due to the time taken for regulatory and ethical approvals for the study at the time of the urgent roll-out of BBIBP-CorV vaccination in Pakistan, we could not unfortunately determine baseline data exactly for the subjects in the BBIBP-CorV vaccine study. However, this study population is similar to the one in which we investigated seroprevalence against SARS-CoV-2 in an uninfected healthcare associated cohort tested prior to vaccinations (conducted March to October 2020). We had observed that seropositivity to spike to be 35% and 21.3% to RBD in our healthy uninfected controls (32). As, many vaccinees here belonged to the larger study we expect that their baseline seroprevalence would be comparable. Neutralizing assays against SARS-CoV-2 were not conducted in this cohort. However, we have shown previously that sera samples positive for IgG to RBD had neutralizing activity against SARS-CoV-2 (73). Therefore, IgG to RBD would be associated with greater protection against disease. We were unable to sample each study subject multiple times during the study period and for some we only have one or two serial samples available.

## Conclusions

Our study provides the important immunological insights as to the effect of inactivated COVID-19 vaccine type, BBIBP-CorV from South Asia. It informs on dynamics and magnitude of antibody responses to vaccination. It also informs on cellular immunity to vaccination and the variability of IFNγ-reactivity to different SARS-CoV-2 antigens, Spike as compared with additional genomic determinants. We observed an age-dependent effect with reduced humoral and cellular responses in those aged 50 years and over, supporting a role for booster vaccinations in this group. The effect of COVID-19 infection immunity after vaccination was evident. It also guides that monitoring vaccination induced immunity to vaccines that include genome wide antigens other than Spike would need to be different from that for Spike mRNA or vector based formulations. Hence, recommendations for COVID-19 vaccinations should be made in the context of local immunity and ongoing transmission in the population.

## Acknowledgements

This study was supported by the Provost’s Academic Priorities Fund, Aga Khan University. Thanks to Paula Alves, iBET/ITQB, NOVA University, Portugal for providing recombinant proteins used in this study. MV was supported by the European Union H2020 ERA project (No 667824 - EXCELLtoINNOV). We thank all the study subjects for participation in this study. Thanks to Ambreen Wasim for assistance with statistical analysis. Also thank Naila Baig Ansari and Nausheen Ahmad for assistance in recruitment of study subjects.

## Contributors

ZH was responsible for conceptualization of the study. ZH, KIM, SHA, KG and EK implemented the study. MV, PA, JPS, EK, SHA, KI, SFM, NTI, KA, MR and ZH guided on methodology. KIM, SQ, SHA, KI, MY and SS performed the experimental work. AH, ZG, SQ, SA, SB, NA, HH, JI assisted with patient recruitment. ZB and JI provided technical resources. MY, SQ, SB, SA, HIM were involved in data curation were involved in data analysis. ZH, KIM, SQ, MV, RH, EK, SHA, KG reviewed and edited the manuscript. All authors have read and approved the final draft of the manuscript.

